# Na/K-ATPase Activity and Ketone Body Metabolism in Long-term Diabetic Rats

**DOI:** 10.1101/567180

**Authors:** Engelbert Buxbaum

## Abstract

The long-term (34 weeks) effect of streptozotocin induced diabetes was assessed in Wistar rats.

Na^+^/K^+^-ATPase activity was measured by ouabain inhibitable ^86^Rb^+^-uptake into erythrocytes. No difference in the rate of Rb^+^-uptake, the K_*m*_ for Rb^+^ or the K_*i*_ for ouabain was detected between normal and diabetic rats. Thus, the change in Na^+^/K^+^-ATPase activity repeatedly described in short-term studies may not translate into a long term physiologically relevant change in ion flux through the sodium pump.

Rats excrete ketone bodies mainly as β-hydroxybutyrate. This compound does not show up with nitroprusside sodium based test sticks, it can however be detected by coupled spectrophotometric assay with hydroxybutyrate dehydrogenase.

Almost half of the diabetic animals reverted to a non-diabetic state during the experiment, followed by at least partial reversal of secondary diabetic damage.

**Abbreviations used:**

PKC: Protein kinase C
DAG: Diacylglycerol
EDTA: Ethylenediamine tetraacetic acid
PIP_3_: Phosphatidylinositol trisphosphate
PBS: Phosphate buffered saline
i.p.: intraperitoneal
HBA: β-hydroxybutyric acid
STZ: streptozotocin
Enzymes: : Na^+^/K^+^-exchanging ATP-phosphohydrolase (Na^+^/K^+^-ATPase), E.C. 7.2.2.13; (R)-3-hydroxybutanoate:NAD^+^ oxidoreductase [1.1.1.30] (β-hydroxybutyrate dehydro-genase)

## 1 Introduction

Diabetes mellitus is a serious disease caused either by the inability to produce insulin (called either type I diabetes, juvenile diabetes or insulin deficiency diabetes) or by the inability of peripheral tissues to respond to insulin (called type II diabetes, adult onset diabetes or insulin resistant diabetes). Much rarer, it may be caused by genetic deficiencies in the insulin signalling pathway (maturity onset diabetes of the young, MODY). Insulin resistance occurs mainly as a result of obesity, and its prevalence is increasing rapidly in developed societies as a result of a diet high in simple carbohydrates and fats, combined with lack of exercise. For Kuwait, this has been described in [1]. For the USA, the Behavioral Risk Factor Surveillance System (BRFSS) offers epidemiological data back to 1985.

Long-term diabetes causes a number of secondary complications in patients like retinopathy, glaucoma, cataract, nephropathy, neuropathy, impaired wound healing and pathologies of the circulatory system. These, much more than diabetes itself, disastrously reduce life expectancy and -quality of the patient.

It has been reported that in diabetic humans as well as laboratory animals the activity of Na^+^/K^+^-ATPase in erythrocytes [2–5], blood vessels [6], peripheral nerves [7–9], brain [10], kidney [11–13], liver [14, 15], pancreas [16], intestine [17], muscle [18, 19], retina and lens [21] is reduced by a factor of up to 7 [12].

Na^+^/K^+^-ATPase uses the chemical energy from the hydrolysis of one molecule of ATP to transport three Na^+^ out of and two K^+^ ions into the cell. Together with K^+^-channels, this causes an electrochemical gradient across the cell membrane which is required for secondary active transport of nutrients and metabolites and for the excitability of neurons and muscle cells. For a review on this enzyme see [22]. Specific inhibitors of Na^+^/K^+^-ATPase are digitalis-like toxins. Of these, ouabain has the highest solubility in water and is hence preferred in the laboratory.

It has been speculated [4] that lower activity of Na^+^/K^+^-ATPase in erythrocytes results in higher osmolarity of the cytosol, resulting in swelling of the erythrocytes. This in turn reduces erythrocyte flexibility, making it more difficult for these cells to pass through narrow capillaries. Thus reduction of Na^+^/K^+^-ATPase activity in erythrocytes could be the molecular cause of microvascular damage in diabetes. If this could be confirmed, erythrocyte Na^+^/K^+^-ATPase activity would be a suitable marker to assess the risk of a patient to develop diabetic complications, it might also be a target for protective drugs. Na^+^/K^+^-ATPase activity reduction could also occur together with other effects of diabetes on cells, rather than being the direct cause of diabetic damage. Even in that case, however, Na^+^/K^+^-ATPase activity could still be a useful diagnostic marker, albeit not a drug target.

Wambach *et al.* [23] have demonstrated that inhibition on Na^+^/K^+^-ATPase (with ouabain) does indeed decrease erythrocyte deformability. Although the potential degree of swelling and reduction in deformability is limited by the filtering activity of the spleen, increased erythrocyte fragility has been demonstrated in blood samples from diabetic patients [24].

Mazzanti *et al.* [25] have shown that differences in Na^+^/K^+^-ATPase activity, membrane fluidity and lipid peroxydation between diabetic and normal humans exist throughout erythrocyte maturation and are superimposed on the changes that occur during erythrocyte aging.

The mechanism of Na^+^/K^+^-ATPase inhibition is unclear, the following causes are discussed:

- Glycation of Na^+^/K^+^-ATPase has been demonstrated in kidney [12] and eye [21] and can change Na^+^/K^+^-ATPase kinetics *in vitro* [26]. However, inhibition is not correlated with blood glucose concentration [3].
- Reduction in cell-membrane phosphatidyl-inositol [7] leads to changes in membrane fluidity and a *reduction* in DAG production, which leads to reduction in protein kinase C activity. However, Yeh *et al.* [27] found that the drug Sorbinil restores Na^+^/K^+^-ATPase activity at a lower concentration than that required for restoration of cell *myo*-inositol levels.
- Hyperglycaemia leads to an *increase* in DAG concentration in the cell and to a stimulation of protein kinase C isoform βII. This is supposed to inhibit Na^+^/K^+^-ATPase directly and via increased prostaglandin production [28].
- Switches in Na^+^/K^+^-ATPase α-subunit isoform expressed on the cell membrane, or an isoform specific shift of Na^+^/K^+^-ATPase molecules from intracellular pools to the plasma membrane [20, 29–31]. Erythrocytes however express essentially only the α1-isoform [32].
- Changes in membrane fluidity by altered lipid metabolism [13, 33].

It should be kept in mind that the route to diabetic damage may vary from tissue to tissue.

Two forms of ATPase-activity reduction have been described:

- Reduction of the number of enzyme molecules per cell (ouabain binding sites, signal in Western blot and mRNA-level) in heart ventricle [31].
- Appearance of substrate inhibition at physiological concentrations of ATP in erythrocytes [3, 34] and lens epithelium [21].

Apart from the mechanism of Na^+^/K^+^-ATPase inhibition in diabetes the physiological significance is also not clear. Does a changed [ATP] *vs* Na^+^/K^+^-ATPase-activity curve *in vitro* actually lead to a significantly reduced ion flux *in vivo*? An easy, well established assay for Na^+^/K^+^-ATPase ion transport activity in erythrocytes is the ouabain inhibitable uptake of ^86^Rb^+^ [35]. Rb^+^ is a congener of K^+^, bound and transported by Na^+^/K^+^-ATPase with similar affinity and efficiency. However, ^86^Rb is handled more easily in the lab than radioactive K-isotopes. A changed Rb-uptake in diabetes has been reported in lens [27], erythrocytes [4] and peripheral nerves [36]. Reduction of Rb-uptake is up to 6-fold [21].

Additionally, ouabain binding kinetics can give an indication of Na^+^/K^+^-ATPase function. Changes in the lipid composition of the membrane lead not only to a change in Na^+^/K^+^-ATPase activity, but also to a changed affinity for this inhibitor [6, 18–20, 33, 34, 37]. This change too can be considerable, an up to 7-fold reduction in K_*i*_ (increase in affinity) has been reported [37].

All these experiments were performed within the first 2 months after induction of diabetes, but secondary complications appear only after several years in human patients. In an attempt to assess the effectiveness of anti-diabetic treatments in preventing long-term complications I decided to look at the erythrocytes of streptozotocinised rats after 8 months, this period in the lifespan of a rat correspond to about 10 years in humans.

Obliteration of β-cells by streptozotocin results in a model of insulin deficiency diabetes. Although this disease is much rarer in humans than insulin resistance diabetes, effects on Na^+^/K^+^-ATPase have been shown to be similar in both diseases [3, 20]. For a review on streptozotocin see [38].

One of the physiological consequences of diabetes is wasting, that is the consumption of body protein and fat. This results in the production of ketone bodies (acetone, acetoacetate and β-hydroxybutyrate), whose concentration is increased in the blood and urine of diabetic patients. Thus apart from the appearance of glucose in urine the appearance of ketone bodies is an easily assessed marker for diabetes. This test is usually performed with test stripes soaked with potassium pentacyanonitrosylferrat (*vulgo*: nitroprusside sodium), which turns cherry-red in the presence of ketones. Surprisingly, I could not find any ketone bodies in urine of diabetic rats using this test, despite clear signs of wasting. I decided to investigate this finding.

## 2 Materials and Methods

Animals were kept in the Animal Resource Centre at the Faculty of Medicine, University of Kuwait. All laws of Kuwait and all rules and regulations of Kuwait University regarding animal handling were strictly observed.

Female Wistar rats, 200–250 g at the beginning of the experiment, were treated with 60 mg/kg streptozotocin (Fluka, 100 mg/ml in 10 mM Na-citrate *p*H 4.5, freshly prepared) i.p. to induce diabetes. Naïve rats were given an equal volume of buffer instead. Rats were kept in colony cages on pine shavings. Food (RM1 9.5 mm pellets, Special Diet Services, Witham, UK) and tab water were provided *ad libitum*, water consumption of each group was measured daily.

Urine glucose and ketone bodies were measured weekly with BMTest7 stripes (Boehringer Mannheim, Germany). Weight was also determined every week.

Blood glucose and total hæmoglobin (B-glucose and hæmoglobin photometers, HemoCue AB, Ängelholm, Sweden) and 24 h urine production were monitored monthly. To avoid unnecessary stress to the animals, tests were performed in the fed state.

In the urine, β-hydroxybutyrate (HBA) was determined by coupled spectrophotometric assay with β-hydroxybutyrate dehydrogenase (obtained as kit from Sigma, St. Louis, USA). It was verified that urine does not interfere with this kit, which is intended for use with blood. Assays were performed as suggested by the manufacturer, except that all volumes were scaled down to 1/10, and a microcuvette was used.

Total urine protein was determined by the method of Lowry *et al.* [39], lysozyme as described in [40].

Animals were sacrificed after 8 months. They were sedated with 400 mg sodium diethylbarbiturate and 300 mg EDTA in PBS i.p., followed by narcosis with 60 mg phenobarbitone (Sagatal, Rhone-Merieux Ltd). This sequence avoids the strong initial excitation seen when phenobarbitone is given alone. The body cavity was opened and a perfusion needle inserted into the aorta from the left ventricle. Rats were perfused with 50 ml PBS, blood was collected from the vena cava.

The collected blood was spun at 350 g for 10 min, supernatant and buffy coat were removed and the erythrocytes washed three times in transport buffer (potassium-free Krebs-Henseleit-buffer [41]: 10 mM Glucose, 1.22 mM NaH_2_PO_4_, 2.50 mM Na_2_HPO_4_, 12 mM NaHCO_3_, 128 mM NaCl, 0.6 mM MgCl_2_, 0.5 mM CaCl_2_) and resuspended in this buffer to 20 % hæmatocrit. The erythrocyte concentration and diameter were determined with a Z2 particle counter (Coulter) after dilution with PBS. Erythrocytes were used on the day of collection to avoid hæmolysis.

To determine ouabain-inhibitable uptake of ^86^Rb^+^, erythrocytes (8 % hæmatocrit), RbCl (0 to 2.5 mM, with NaCl added to keep the osmolarity constant) and ouabain (0 to 50 µM) were pre-incubated in transport buffer for 30 min at 37 ^?^C. The reaction was started by addition of 29 kBq ^86^RbCl (Amersham, 11 µM final concentration). Final volume was 500 µL. Samples of 50 µL were withdrawn every 15 min and placed onto gradients of 200 µL dibutylphtalate over 100 µL 3 M KOH. Erythrocytes were spun immediately through the dibutylphtalate for 30 s at 10000 rpm in an Eppendorf centrifuge, the aqueous phase and most of the dibutylphtalate were removed by aspiration. The Cherenkov-radiation in the KOH-phase was determined in a β-counter (Beckman), using ^3^H-settings. Background was calculated from the [ouabain]-dependence of the count rate at a fixed Rb-concentration.

## 3 Results

### 3.1 Diabetic state of the animals

In total 21 animals were treated with streptozotocin, 5 did not become diabetic in the first place, a further 7 animals reverted to a non-diabetic state later. These animals were combined into a separate group, called revertants. 9 animals stayed diabetic throughout the 8 months of the experiment. Prandial blood glucose levels in these animals were in excess of 22 mM, the maximum range of the HemoCue blood glucose photometer, urine glucose was 4+ on the test stripes.

The amount of urine produced increased from (0.054 ± 0.025) (*n* = 21) in normal to (0.383 ± 0.172) mL/(g * d) (*n* = 9, max = 0.782 mL/(g * d)) in diabetic rats. Nevertheless, these animals stayed active and inquisitive, even assertive, with normal grooming and social behaviour.

It is often said that diabetic rats cannot be maintained for extended periods. In my experience, however, the only additional measures needed to maintain these animals were to ensure an adequate water supply (especially over the weekend) and litter changes every other day. Because of wasting, experimental and control animals were always kept in separate cages.

Dissection showed wasting of muscle and non-essential organs, for example of uterus and ovaries (fig. 1).

**Figure 1:**
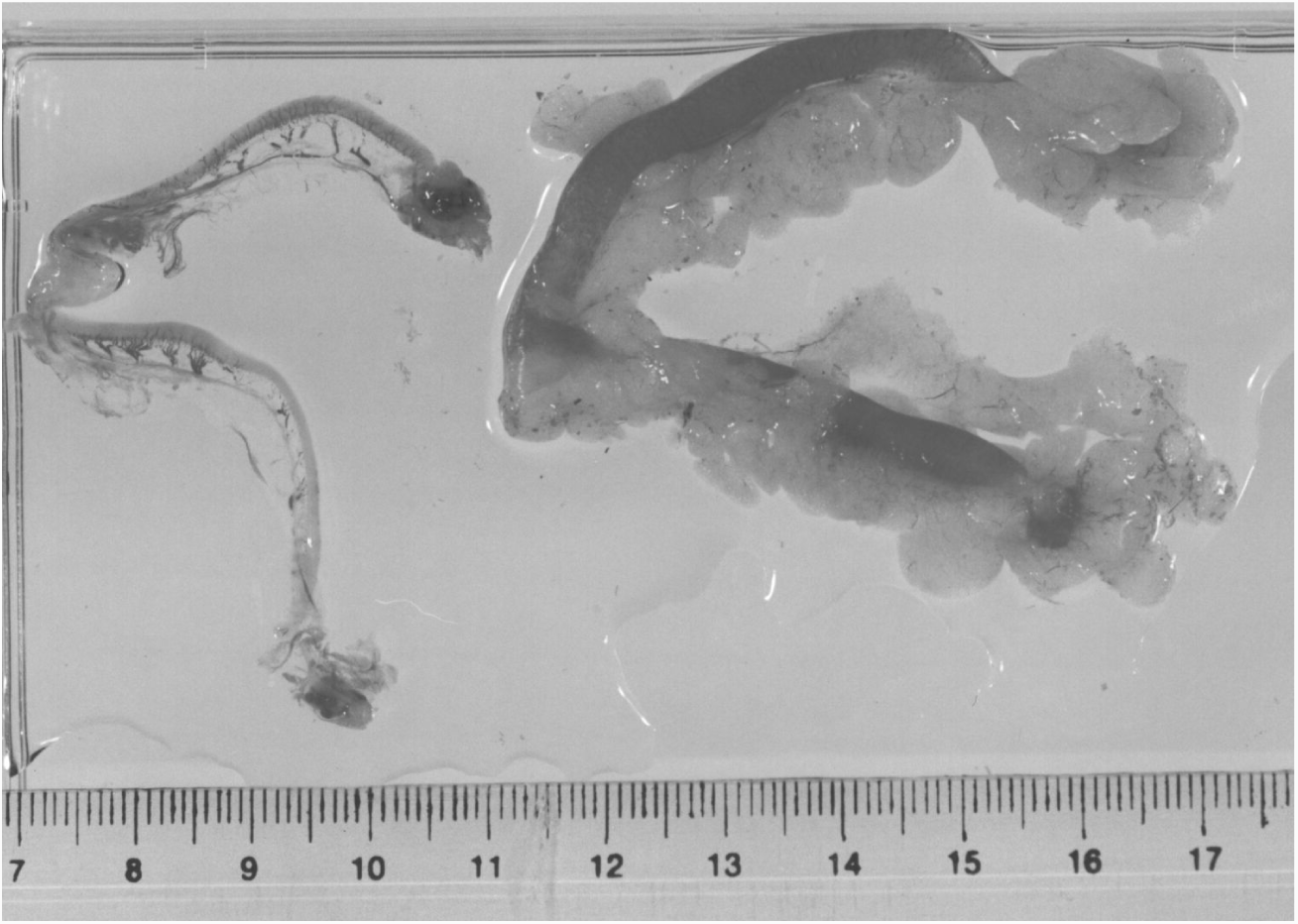
Ovaries and uterus from a normal (*right*) and a diabetic rat (*left*). Severe wasting can be observed in case of the diabetic rat, the organs are much smaller and lack adipose tissue. Scale is in cm.

Protein secretion in diabetic animals was increased both for large proteins (showing damage to the glomerulus) and for small proteins like lysozyme (showing reduced reuptake in the tubuli). In this respect, the revertants were a mixed group, some showing values reminiscent of naïve, others of diabetic animals. This may not be surprising however, as the reversion to a non-diabetic state occurred at different times in different animals.

**Table 1:** Clinical data 8 months after injection of STZ (arithmetic means and standard deviation). Statistical significance was determined in a 2-sided t-test with the naïve group and is marked (-) not significant, (*) significant (P_0_ *<* 5%) and (**) highly significant (P_0_ *<* 0.1%).

| Parameter | Unit | naïve | revertants |  | diabetics |  |
| --- | --- | --- | --- | --- | --- | --- |
| total haemoglobin | g/dL | $15.8 \pm 0.9$ | $14.8 \pm 1.3$ | - | $6.5 \pm 0.9$ | - |
| Blood Glucose | mM | $9.0 \pm 2.5$ | $10.5 \pm 2.3$ | - | $>22$ | ** |
| Weight | g | $358 \pm 21$ | $291 \pm 34$ | - | $263 \pm 28$ | * |
| Urine Glucose | 0-4 | $0 \pm 0$ | $0 \pm 0$ | - | $4 \pm 0$ | * |
| Urine Volume | mL/d | $14.2 \pm 4.6$ | $18 \pm 7$ | - | $101 \pm 28$ | * |
| Urine Protein | mg/d | $9.6 \pm 2.1$ | $9.6 \pm 2.9$ | - | $137 \pm 40$ | * |
| Urine Lysozyme | $\mu\text{g/d}$ | $60 \pm 27$ | $135 \pm 147$ | - | $369 \pm 102$ | ** |
| Urine HBA | mg/dL | $1.8 \pm 0.8$ | | | $1.58 \pm 0.57$ | - |
| Urine HBA | mg/d | $26 \pm 16$ | | | $159 \pm 65$ | ** |
| Erythrocyte ∅ | $\mu\text{m}$ | $6.14 \pm 0.44$ | $5.78 \pm 0.17$ | - | $5.7 \pm 0.8$ | - |
| $K_m$ Rb | mM | $0.83 \pm 0.12$ | $0.91 \pm 0.38$ | - | $1.45 \pm 0.89$ | - |
| $K_i$ Ouabain | $\mu\text{M}$ | $21 \pm 33$ | $31 \pm 23$ | - | $21 \pm 13$ | - |
| Rb Transport rate | fmol/(min * Ery) | $4.6 \pm 0.8$ | $5.2 \pm 4.2$ | - | $6.1 \pm 3.9$ | - |

Total hæmoglobin was marginally, but not significantly increased in diabetic animals, an effect usually attributed to changes in oxygen affinity following problems with 2,3-bisphosphoglycerate metabolism.

### 3.2 Ketone body metabolism

Since the diabetic animals showed clear signs of wasting, it was expected to find ketone bodies in their urine as product of fatty acid and protein catabolism. However, no ketone bodies could be found using the BMtest7 sticks at any of the weekly testings. One possible explanation was that rats excrete mostly β-hydroxybutyric acid (HBA), which does not contain a keto-group and thus does not react with the nitroprusside sodium contained in the test strips. Indeed, with a coupled spectrophotometric assay HBA could be detected, but in concentrations not significantly different from naïve animals. However, diabetic rats produce a very large urine volume, thus the daily excretion of HBA is 6 times higher in diabetic than in normal rats (159 vs 26 mg/d).

### 3.3 Na^+^/K^+^-ATPase activity

Contrary to what might have been expected from the literature, neither the maximal rate of Rb^+^-uptake, nor the K_*m*_ for Rb^+^ changed as a consequence of increased blood glucose levels in diabetic rats, the affinity for ouabain did not change either. No changes in erythrocyte diameter were observed (fig. 3).

## 4 Discussion

### 4.1 Ketone body metabolism

Just like humans, rats seem to produce mostly β-hydroxybutyrate and little acetoacetate or acetone. Since HBA does not contain a keto-group, it does not show up on urine test stripes containing nitroprusside sodium, a well-known clinical problem in humans.

Although the HBA-concentration in the urine of diabetic rats is similar to that of naïve controls, this concentration needs to be multiplied by the 7-fold higher urine volume to compare daily excretion. This experiment makes a nice case study in a course on clinical biochemistry, showing how the physiological situation of a patient needs to be taken into account when interpreting laboratory results.

### 4.2 Na^+^/K^+^-ATPase activity

The results of this study do not support the hypothesis that microvascular damage in diabetic subjects is caused by a reduction of erythrocyte Na^+^/K^+^-ATPase activity, followed by swelling of the erythrocytes [4], at least in this simple form.

No changes of erythrocyte diameter could be detected in rats eight months after obliteration of islet β-cells with streptozotocin. This result however needs to be viewed with great caution, the complex shape of red cells may change considerably before a particle counter would notice a change in diameter.

More importantly, no changes were observed in the rate of ^86^Rb^+^-uptake, the K_*m*_ for Rb^+^ and the K_*i*_ for ouabain. The results for the diabetic rats fall neatly into the range observed for both naïve animals and revertants, despite the severe diabetes maintained for eight months.

It would have been interesting to do further determinations, like ouabain binding capacity, Na^+^/K^+^-ATPase activity at physiological ATP-concentrations and an activity vs [ATP]-curve. However, with the limited amount of material available these further investigations were not feasible.

Changes in Na^+^/K^+^-ATPase activity and response to ATP-concentrations in diabetes are now well documented [2–11, 13–17, 21, 27, 36] in several tissues (including erythrocytes) and physiologically relevant changes in Rb-uptake have been demonstrated in lens, nerve and erythrocytes [4, 27, 36], but only in the short term (several days to 2 months). Changes in ouabain affinity have also been described [6, 18–20, 33, 34, 37]

The data reported here have to be compared with the data reported in the literature: a 7-fold change in the rate of ATP-hydrolysis [12], an up to 27-fold decrease in the affinity for ATP [12], a 7-fold change in ouabain K_*i*_ [37] and a 6-fold change in potassium flux [21].

**Figure 2:**
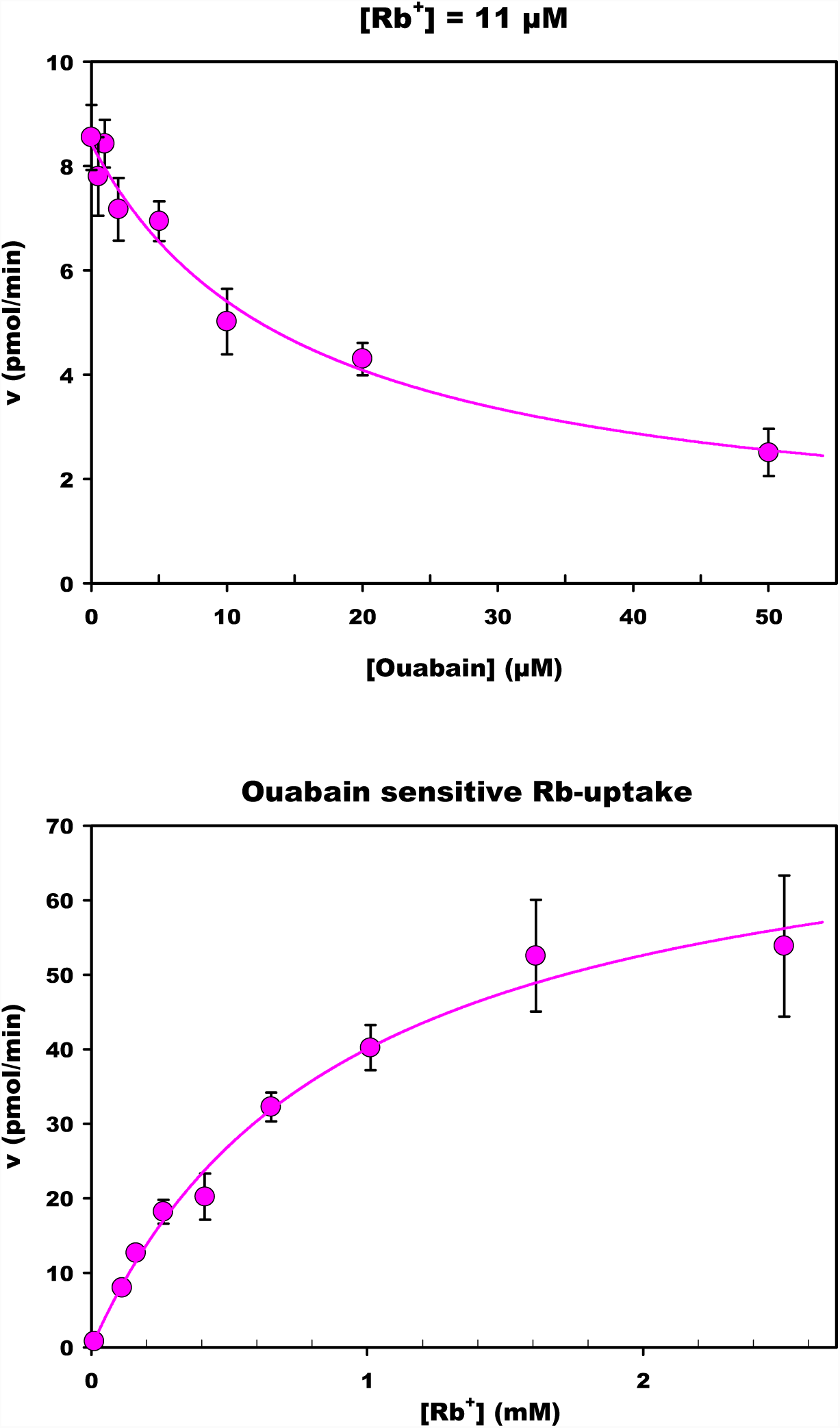
Original Data for Rat 2 (naïve). *Top:* Inhibition of Rb-uptake by different ouabain concentrations. *Bottom:* Effect of the Rb-concentration on the rate of Rb-uptake. The fitted curves have the equations 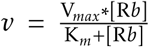 and 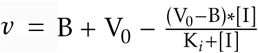 respectively (with B the background and [I] the ouabain concentration). Note that the kinetics of Na^+^/K^+^-ATPase interaction with K^+^ and Rb^+^ is known to be described by a Henri-Michaelis-Menten equation despite the fact that two ions are bound [42–45].

**Figure 3:**
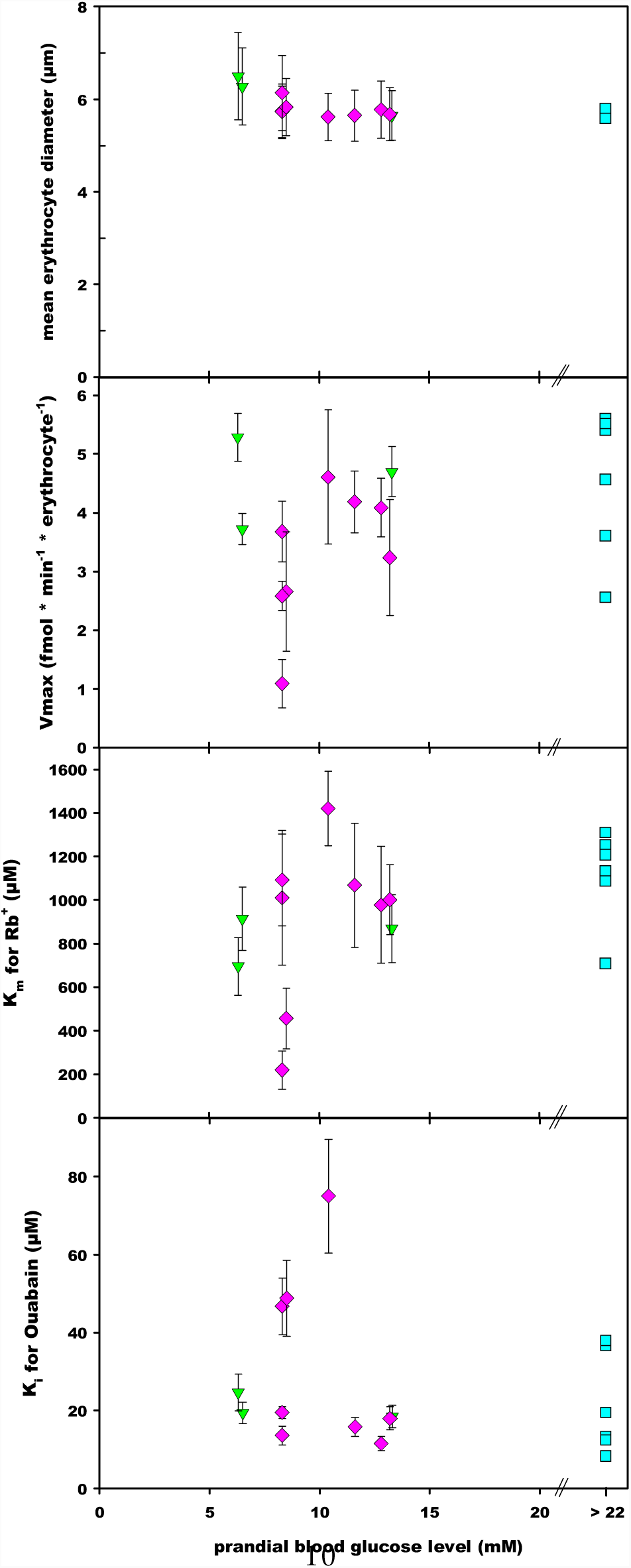
Effect of diabetes on rat erythrocytes. *Top:* mean erythrocyte diameter. *2nd:* maximum rate of Rb-transport. *3rd:* K_*m*_ for Rb. *bottom:* K_*i*_ for ouabain. Error bars not shown for diabetic rats to improve clarity. Triangle: naïve, diamond: revertant, square: diabetic.

The interpretation of these studies and our own is made more complicated by the fact that Na^+^/K^+^-ATPase activity shows high individual variability from rat to rat. Simulations (not shown) indicate that in this study changes by a factor of 2–3 would have been detected as significant, half than what was reported in other studies.

Changes in Na^+^/K^+^-ATPase activity appear to be a fast response to perturbations of glucose and polyol metabolism, to which at least the rat can slowly adapt. The longer time period used here (34 weeks) seems warranted in view of the slow appearance of diabetic damage in human patients.

This conclusion is supported by other results in the literature:

- The number of Na^+^/K^+^-ATPase molecules in rat heart ventricle is decreased two weeks after induction of diabetes, but begins to recover after four, further data points are unfortunately not available [31].
- Exposure of rabbit aortic endothelium to high glucose medium *in vitro* resulted in a dose dependent, saturatable inhibition of ouabain-inhibitable Rb-uptake (i.e. Na^+^/K^+^-ATPase activity) within 60 min [46]. Like the Na^+^/K^+^-ATPase inhibition observed *in vivo*, the effect was prevented by *myo*-inositol or Sorbinil.

The speed of Na^+^/K^+^-ATPase inhibition may in part explain the need for constant effective control of blood glucose concentration in human diabetics. Clinical studies show that even relatively short term lapses appear to contribute to long term diabetic damage. The sequence of events however appears to be more complicated than that suggested by Kowluru *et al.* [4]. Caution is therefore advised when Na^+^/K^+^-ATPase activity is used as marker in studies of possible pharmaceuticals against secondary complications of diabetes [7, 8, 13, 14, 27, 35, 47] or in other experimental situations [17, 18, 33].

### 4.3 Reversibility of STZ-diabetes and secondary diabetic complications

One interesting finding of this study was that given enough time many rats can overcome the damage done by STZ and start producing insulin again. Once normoglycæmia is restored, secondary damage, for example to the kidney, seems to be repairable. Both glomerular and tubuli function returned to normal in at least some of the revertant rats, as determined by protein and lysozyme excretion. General health, for example body weight, also improved slowly in revertants. Such slow recovery for example of kidney function is also observed in human patients after pancreas transplantation [48].

Of course STZ toxicity is a short term event, and re-establishment of a viable population of β-cells is possible. In human type 1 diabetes this is prevented by a continuing autoimmune reaction. Could this autoimmune reaction be controlled to allow self-healing to take place in human patients?

## Acknowledgements

This work was supported by Kuwait University. The help of Mona Bakir, Mona N. Al-Rustom and T.S. Srikumar is gratefully acknowledged. Thanks are also due to the staff of the Animal Resource Centre of Kuwait University for maintaining the rats. I am indebted to Drs F.M. Al-Awadi, C. Cojocel and W. Renno for their valuable input into this study.

